# Dryland cropping system, weed communities, and disease status modulate the effect of climate conditions on wheat soil bacterial communities

**DOI:** 10.1101/2020.04.13.040253

**Authors:** Suzanne Lynn Ishaq, Tim Seipel, Carl Yeoman, Fabian D. Menalled

**Author notes:** Corresponding authors: Suzanne Ishaq, University of Maine, School of Food and Agriculture, 5735 Hitchner Hall, Room 130A, Orono, ME 04469, Fabian Menalled, Montana State University, Department of Land Resources and Environmental Sciences, 720 Leon Johnson Hall, Bozeman MT, 59717. University of Maine, School of Food and Agriculture, Orono, ME 04469. Emails: SI, TS, CJY, FM.

## Abstract

Little knowledge exists on whether soil bacteria are impacted by cropping systems and disease status in current and predicted climate scenarios. We assessed the impact of soil moisture and temperature, weed communities, and disease status on soil bacterial communities across three cropping systems: conventional no-till (CNT) utilizing synthetic pesticides and herbicides, 2) USDA-certified tilled organic (OT), and 3) USDA-certified organic with sheep grazing (OG). Sampling date within the growing season, and associated soil temperature and moisture, exerted the greatest effect on bacterial communities, followed by cropping system, Wheat streak mosaic virus (WSMV) infection status, and weed community. Soil temperature was negatively associated with bacterial richness and evenness, while soil moisture was positively associated with bacterial richness and evenness. Both soil temperature and moisture altered soil bacterial community similarity. Inoculation with WSMV altered community similarity, and there was a date x virus interaction on bacterial richness in CNT and OT systems, as well as an interaction between WSMV x climate. In May and July, cropping system altered the effect of climate change on the bacterial community composition in hotter, and hotter and drier conditions not treated with WSMV, as compared to ambient conditions. In areas treated with WSMV, there were interactions between cropping system, sampling date, and climate conditions, indicating the effect of multiple stressors on bacterial communities in soil. Overall, this study indicates that predicted climate modifications as well as biological stressors play a fundamental role in the impact of cropping systems on soil bacterial communities.

**IMPORTANCE:** Climate change is affecting global moisture and temperature patterns and its impacts are predicted to worsen over time, posing progressively larger threats on food production. In the Northern Great Plains of the United States, climate change is forecasted to increase temperature and decrease precipitation during the summer, and is expected to negatively affect cereal crop production and pest management. In this study, temperature, soil moisture, weed communities, and disease status had interactive effects with cropping system on bacterial communities. As local climates continue to shift, so too will the dynamics of above- and below-ground associated bio-diversity which will impact food production and the need for more sustainable practices.

## INTRODUCTION

Climate change affects soil moisture content and temperature which, in turn, impacts: crop production and nutritional value (1–4); pest abundance, dynamics, and management (4–7); as well as overall ecosystem resiliency (8). Determining how climate change modifies multitrophic interactions between crops, weeds, pathogens, and soil microbial communities is complex (9), yet critical, as crop production relies on healthy soil and microbially-mediated nutrient cycling (10, 11). Microbial α-diversity in soil is linked to plant growth stage (12). Low microbial α-diversity in soil is associated with impeded plant growth, and early senescence of *Arabidopsis thaliana* (13). With the knowledge that climate change will fundamentally change the dynamics of agricultural ecosystems, we must increase our understanding of the mechanisms driving biological and environment stress to secure the sustainability of agricultural production, (1, 14, 15).

The Northern Great Plains of the United States is a major global cereal-producing region where the effects of climate change are already being felt (16, 17). Over the next 30 years, mean temperature is predicted to increase by 2.5 – 3.3°C in this region (17, 18). Soil microbial community structure and function may be altered due to their temperature sensitivity (19–21). Coupled with predicted decreases in summer precipitation, hotter and drier conditions during the growing season will result in crop stress (17) which has the potential to further alter soil microbial communities. In periods of drought, microbial diversity is reduced (22, 23) as is their ability to cycle soil nitrogen (24). Drought can also cause plants to prioritize relationships with fungi over bacteria, reducing the transfer of nutrients and contributing to the crash of the bacterial community (25, 26). Further, as climate change alters the composition of plant communities and their nutrient content (27, 28), the composition of plant liter and residues is altered. This change in soil inputs, in turn, modifies plant-microbe relationships (29–31) and reduces the available nutrients recycled into soil (22, 30).

Climate change is also predicted to worsen the effects of plant pathogens, including *Wheat streak mosaic virus* (WSMV; genus *Tritimovirus*), either by altering the dynamics of vector transfer or by decreased plant resistance to infection (7, 32). WSMV is transmitted by wheat curl mites (*Aceria tosichella*), occurs across the North American Great Plains, and can make plants more susceptible to the effects of climate change by hindering root development and water uptake (33). To our knowledge, no study has formally assessed the potential link between WSMV infection and plant-, rhizosphere-, or root-associated microbial communities. While un-explored, it is possible that the WSMV viral infections which alters root structure or function, and thus the capacity for plants to interact with soil microbiota.

In industrial (contemporarily referred to as conventional) cropping systems, management approaches focused on maximizing production are based on regular applications of synthetic inputs in the form of fertilizers and pesticides (34). In recent years, shifted consumer demands and new market opportunities have developed organic production into a major agricultural, economic, and cultural force (35, 36). However, organic cropping systems rely heavily on tillage for weed management and cover-crop termination. Due to the negative consequences that tilling has on the physical, chemical, and biological properties of soils in the semi-arid ecosystems that dominate large sections of the Northern Great Plains, there is a growing interest among farmers and researchers to reduce soil disturbance practices in organic systems (37–39). In this context, the integration of crop and livestock production has been proposed as a sustainable approach to terminate cover crops, manage crop residues, and control weeds while reduce tillage intensity (40–42), yet very few studies exist on the impact of integrated livestock management on soil quality or microbial communities (23) or disease resistance.

Differences among cropping systems affect plant communities, including species’ abundance, composition, and growth (43, 44) which, in turn, modifies microbial communities in the rhizosphere (23, 45, 46). Although previous studies have evaluated the role of microbial communities in crop yields and crop-weed competition (47), fewer explore the extent to which root-associated bacteria are impacted by cropping systems, weeds, and plant disease in current and predicted climate scenarios. The aim of our study was to assess changes in soil bacterial communities due to warmer and drier climate conditions and the presence of WSMV across contrasting cropping systems and their associated weed communities. We hypothesized that: 1) bacterial community richness and evenness would be reduced by climate or WSMV infection; 2) cropping systems that promote bacterial richness would be more resistant to alterations from climate and WSMV infection; and 3) more diverse bacterial communities would have a more stable bacterial community membership over the growing season and in response to increased soil temperature, decreased precipitation, and WSMV.

## RESULTS

### Bacterial diversity and evenness

Soil temperature during the growing season (Fig S1) was a strong driver of bacterial species’ richness (Table 1); with fewer bacterial OTUs (97% cutoff) observed in soil during hotter temperatures (Fig 1A). Increased soil temperature reduced the evenness of bacterial species’ (Table 1). Hotter soil temperatures were negatively associated with the presence or relative abundance of bacterial taxa that were significantly important features in the model (Figure 1B). The most abundant of those taxa included members of *Blastococcus*, Bacillales, Micromonosporaceae, Intrasporangiaceae, *Sphingomonas*, Microbactericeae, and *Streptomyces* (Fig 1B).

**Table 1.** Effect of treatment factors and their interactions on observed soil bacterial richness and evenness. Richness is measured as bacterial taxa counts and evenness of taxa abundance on a scale from 0 to 1 (each species having equal abundance). Comparisons were made using a linear mixed effects model accounting for repeated measures of subplots within replicated blocks and significance was determined via Type III ANOVA with Satterthwaite’s approximation.

|  | Observed Richness |  |  | Shannon Evenness |  |  |
| --- | --- | --- | --- | --- | --- | --- |
|  | Sum Sq | F value | P value | Sum Sq | F value | P value |
| Cropping System (C) | 275,240 | 4.53 | 0.012 | 0.051 | 6.28 | 0.002 |
| Soil Temperature (T) | 2,513,973 | 82.82 | <0.001 | 0.080 | 19.64 | <0.001 |
| Soil Moisture (M) | 1,201,709 | 39.59 | <0.001 | 0.028 | 6.97 | 0.009 |
| WSMV (V) | 56,697 | 1.87 | 0.173 | 0.001 | 0.14 | 0.708 |
| Date* | 2,203,758 | 28.71 | <0.001 | 0.104 | 8.6 | <0.001 |
| C:T | 442,913 | 7.30 | 0.001 | 0.072 | 8.84 | <0.001 |
| C:M | 217,218 | 3.58 | 0.029 | 0.015 | 1.89 | 0.154 |
| T:M | 845,614 | 27.86 | <0.001 | 0.025 | 6.07 | 0.014 |
| C:V | 110,178 | 1.81 | 0.165 | 0.001 | 0.17 | 0.844 |
| T:V | 49,186 | 1.62 | 0.204 | 0.001 | 0.32 | 0.571 |
| M:V | 23,029 | 0.76 | 0.385 | 0.001 | 0.17 | 0.683 |
| C:T:M | 179,405 | 2.96 | 0.054 | 0.010 | 1.24 | 0.293 |
| C:T:V | 59,576 | 0.98 | 0.376 | 0.008 | 0.96 | 0.385 |
| C:M:V | 11,219 | 0.18 | 0.831 | 0.002 | 0.21 | 0.812 |
| T:M:V | 4,413 | 0.15 | 0.703 | 0.001 | 0.37 | 0.546 |
| C:T:M:V | 8.038 | 0.13 | 0.876 | 0.005 | 0.59 | 0.556 |

\*Factor used in simple model.

**Fig 1.**
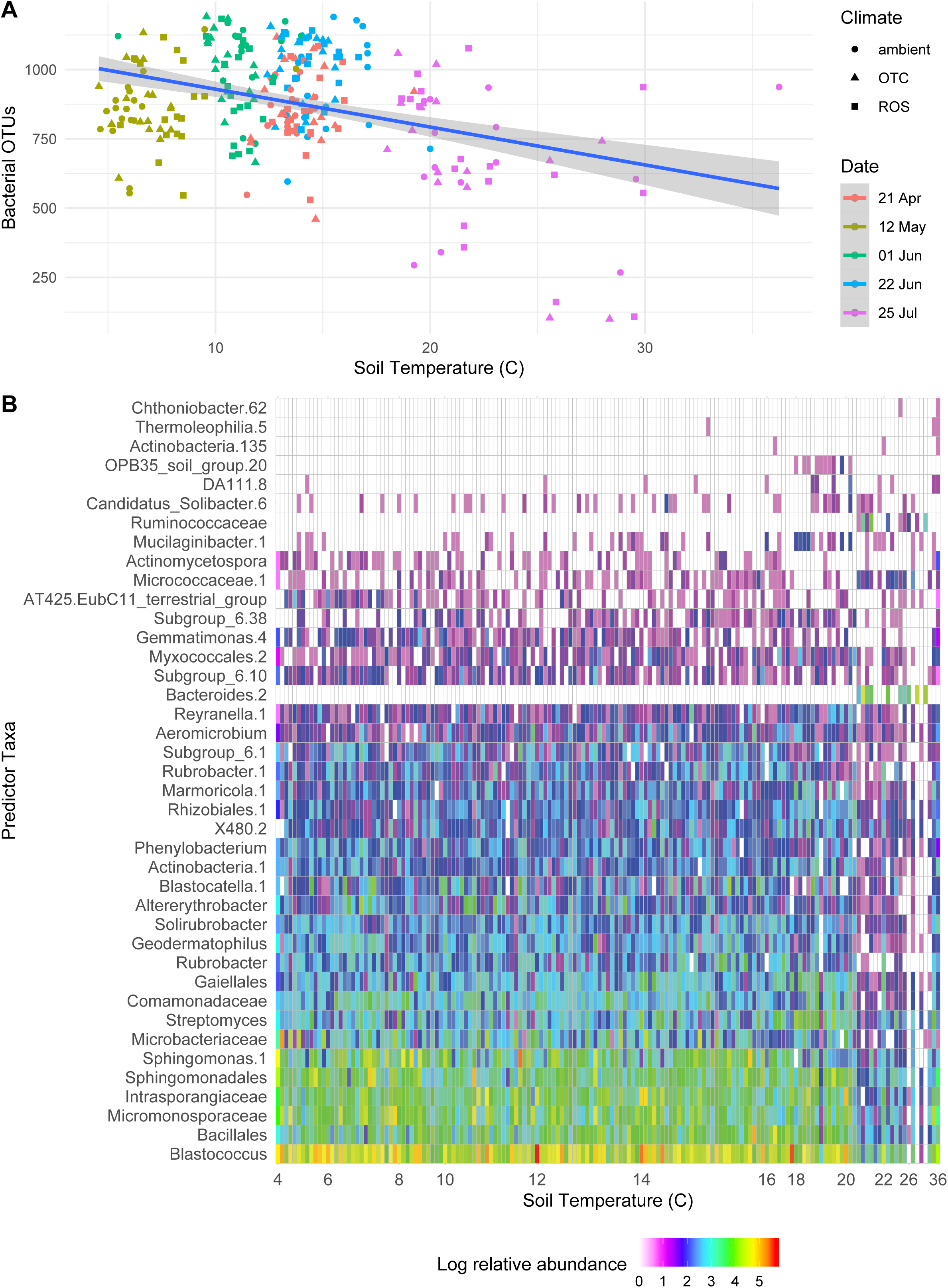
Effect of soil temperature on soil bacterial communities. A) Soil temperature was negatively correlated with soil bacterial richness. B) Relative abundance of soil bacterial by temperature over the 2016 growing season, selected as important features by random forest classification. Taxa are arranged by total relative abundance, and only statistically significant taxa features are shown. Model explained 45% of variance.

Soil moisture during the growing season (Fig S2) positively impacted total bacterial species’ richness (Fig 2A, Table 1) and evenness (Table 1), though not as strongly as temperature did. Across all samples, soil temperature and soil moisture were not correlated with each other (lmer, *p* > 0.05; Fig S3). Soil moisture impacted the relative abundance of bacterial species in different ways (Fig 2B). For example, *Aeromicrobium* were more abundant at low soil moisture levels, *Sphingomonas* were more abundant at high moisture, and *Phenylobacterium* were most abundant at moderate levels of soil moisture (Fig 2B).

**Fig 2.**
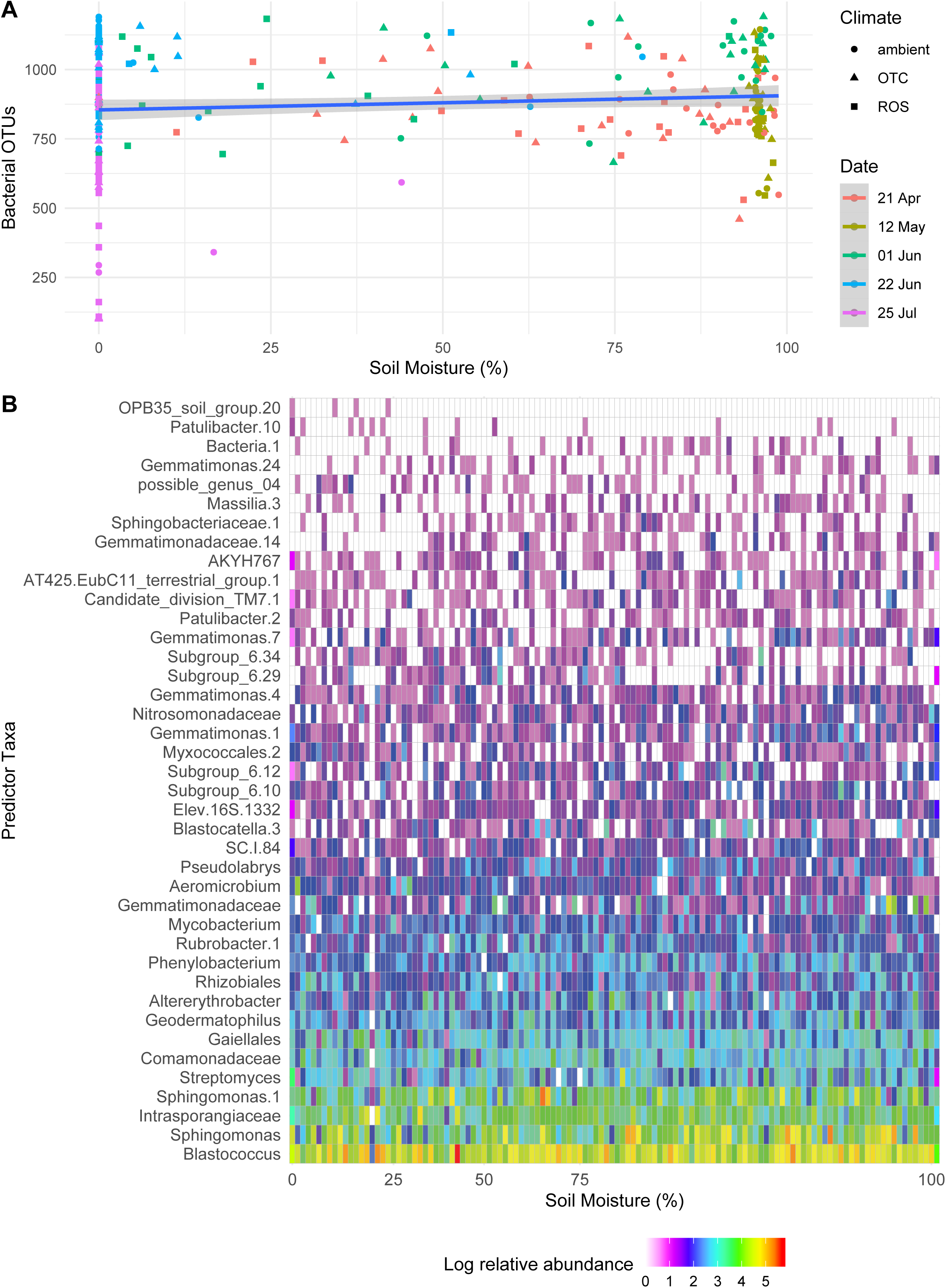
Effect of soil moisture on soil bacterial communities. A) Soil moisture (% of saturation) was positively correlated with soil bacterial richness. B) Relative abundance of rhizosphere bacteria affected by soil moisture from all subplots across the 2016 growing season, selected as important features by random forest classification (*p* < 0.05). Model explained 32% of variance.

Cropping system interacted with climate treatment to affect bacterial richness and evenness (Fig 3; Table 1). Bacterial richness at ambient conditions peaked in early June for all three cropping systems, while richness in both hotter, and hotter and drier conditions peaked in late June in the CNT and OG systems (Fig 3A). Bacterial richness in OG subplots was affected by soil temperature (lmer, Estimate = 35, F = 2.997, *p* = 0.003), and moisture (lmer, Estimate = 6, F = 2.203, *p* = 0.03).

**Fig 3.**
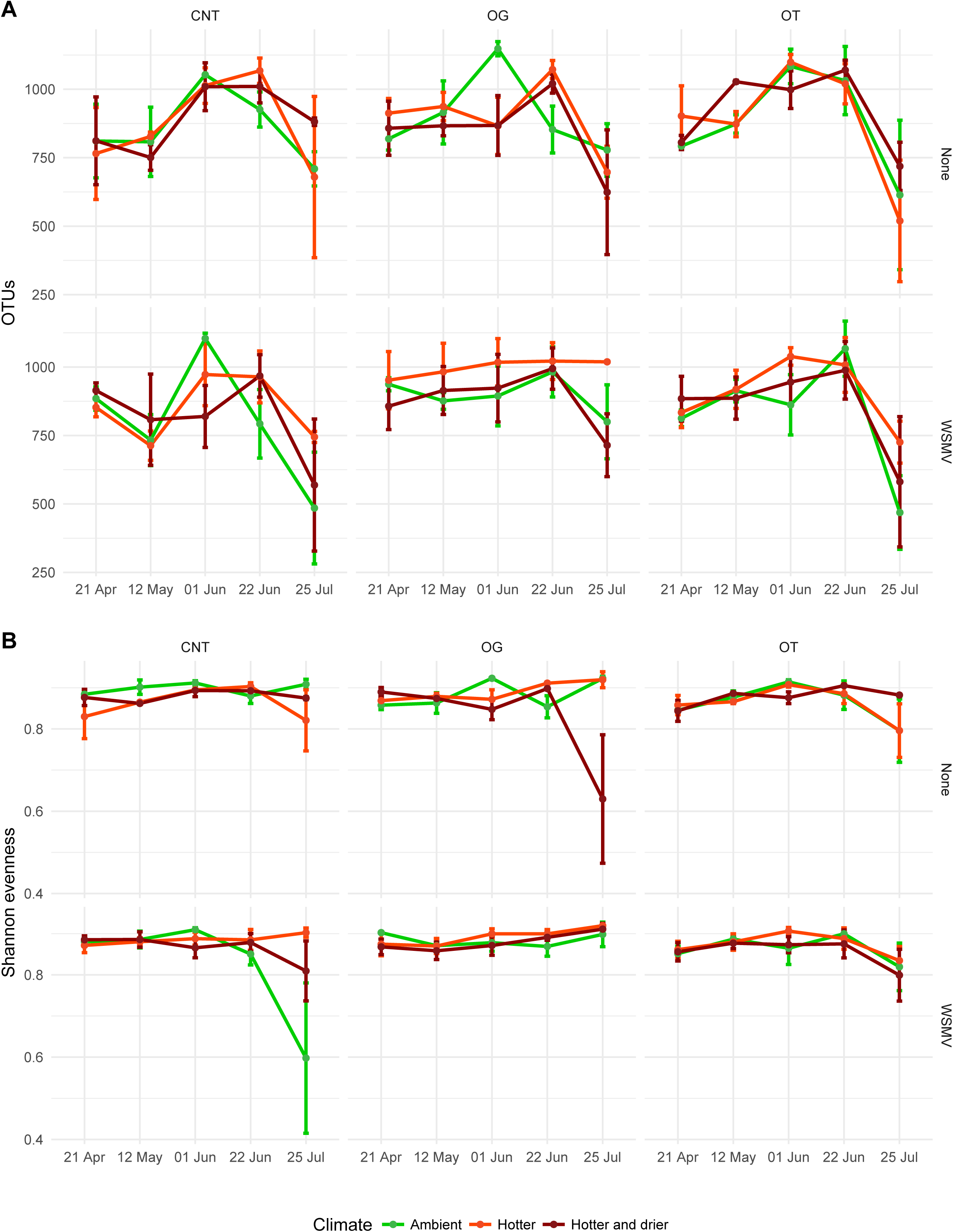
Soil bacterial richness and evenness over the 2016 growing season. A) species-level richness and B) species-level evenness by cropping system (conventional no-till, CNT; organic grazed, OG; organic tilled, OT), climate conditions (ambient, hotter, hotter and drier), and pathogen infection (Wheat streak mosaic virus, WSMV; no WSMV, none). Error bars show Standard Error of Means (SEM).

Inoculation with WSMV resulted in 6 CNT positive samples with mean infection rate within subplots of 4.4%, 9 OG positives with mean infection rate of 13.3%, and 7 OT positives with mean infection rate of 3.2% (Table S1). Overall, inoculation with WSMV had no effect on bacterial species’ richness or evenness (Fig 3; Table 1). When comparing CNT and OT subplots, there was a date x virus interaction on bacterial richness (lmer, F = 2.6792, *p* = 0.039). OT subplots at the end of July that had been inoculated with WSMV showed reduced bacterial richness (lmer, F = 2.019, *p* = 0.046), as did all hotter treatments in July treated with WSMV (F = 3.046, *p* = 0.003) and hotter OG treatments inoculated with WSMV in April (F = 2.039, *p* = 0.044), May (F = 2.088 *p* = 0.039), and late June (F = 2.192, *p* = 0.03). Weed species’ diversity, and percent coverage or biomass, did not alter bacterial richness across all subplots (lm, *p* > 0.05).

### Bacterial community stability

Soil temperature impacted bacterial community similarity (Table 2). Hotter soil temperatures were associated with increased variation in bacterial community heterogeneity and dispersion (betadisper, F = 3.3579, *p* < 0.001), i.e. in warmer temperatures the bacterial communities were more dissimilar across and within a treatment group. Soil moisture also altered soil bacterial community similarity (Table 2), but did not affect the amount of variation (heterogeneity) within bacterial communities (betadisper, *p* > 0.05). Soil moisture did not have an effect on homogeneity when considering healthy and WSMV subplots separately, to account for the effect of WSMV on plants’ abilities to uptake water. Soil bacterial community similarity was impacted by the interaction of cropping system and climate (Table 2).

**Table 2.** PERMANOVA of treatment factors and their interactions on soil bacterial communities. Comparisons were made accounting for repeated measures of subplots and with replicate blocks as a stratification. Significance was determined as: < 0.001 = ***, 0.001 – 0.009 = **, 0.01 – 0.05 = *, 0.05 - 0.1 = t (trending).

|  | Bray-Curtis |  |  |  |
| --- | --- | --- | --- | --- |
|  | F | R<sup>2</sup> | P value |  |
| Cropping system (C) | 0.88 | 0.007 | 0.001 | *** |
| Soil Moisture (M) | 3.67 | 0.014 | 0.001 | *** |
| Soil Temperature (T) | 2.00 | 0.007 | 0.004 | ** |
| Virus (V) | 1.63 | 0.006 | 0.001 | *** |
| Date* | 5.27 | 0.077 | <0.001 | *** |
| C:M | 1.15 | 0.008 | 0.213 |  |
| C:T | 1.69 | 0.012 | 0.005 | ** |
| M:T | 2.88 | 0.011 | 0.001 | *** |
| C:V | 1.00 | 0.007 | 0.189 |  |
| M:V | 1.82 | 0.007 | 0.015 | * |
| T:V | 2.93 | 0.011 | 0.001 | *** |
| C:M:T | 1.03 | 0.008 | 0.411 |  |
| C:M:V | 1.56 | 0.011 | 0.009 | ** |
| C:T:V | 1.25 | 0.009 | 0.107 |  |
| M:T:V | 2.22 | 0.008 | 0.001 | *** |
| C:M:T:V | 1.25 | 0.009 | 0.093 | t |

\*Factor used in simple model.

Here, stability (interpreted as no significant difference in bacterial **β-**diversity) was similar between ambient and treatment subplots over time. Lower OTU richness correlated with a higher similarity between ambient and manipulated hotter and drier subplots (lm, F = 1291, p < 0.001), and this was most evident early and late in the growing season (Fig 4). The temporal stability of a bacterial community against climate change was not associated with a lower fold difference in OTUs between ambient and hotter, and ambient and hotter and drier subplots (Fig 5). While the most stable soil communities did have more bacterial OTUs, a loss of OTUs was not necessarily associated with having lower community similarity (Fig 5).

**Fig 4.**
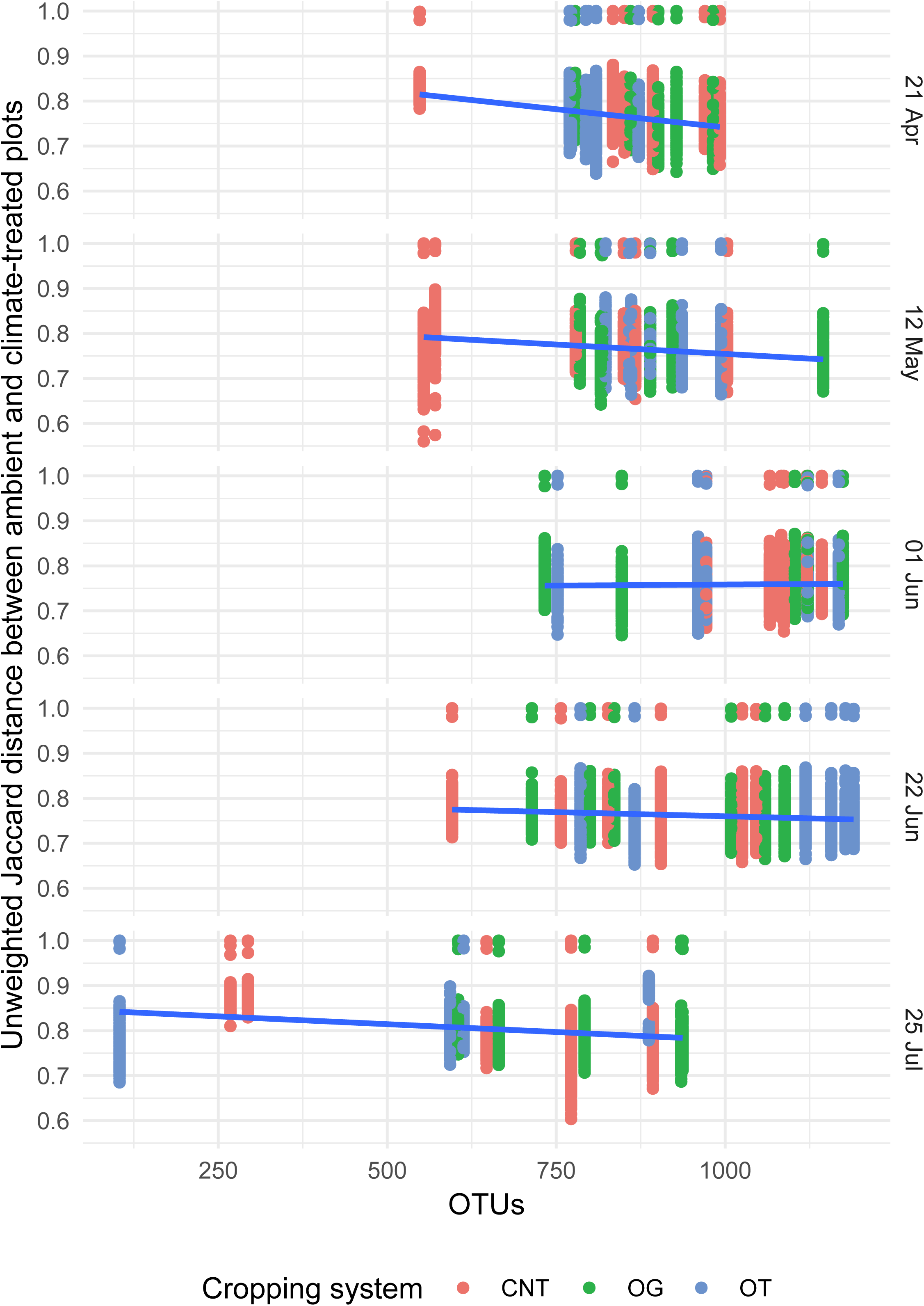
Soil bacterial community similarity between ambient and climate-treated subplots correlation with bacterial OTUs. Cropping systems include conventional no-till (CNT), organic grazed (OG), and organic tilled (OT).

**Fig 5.**
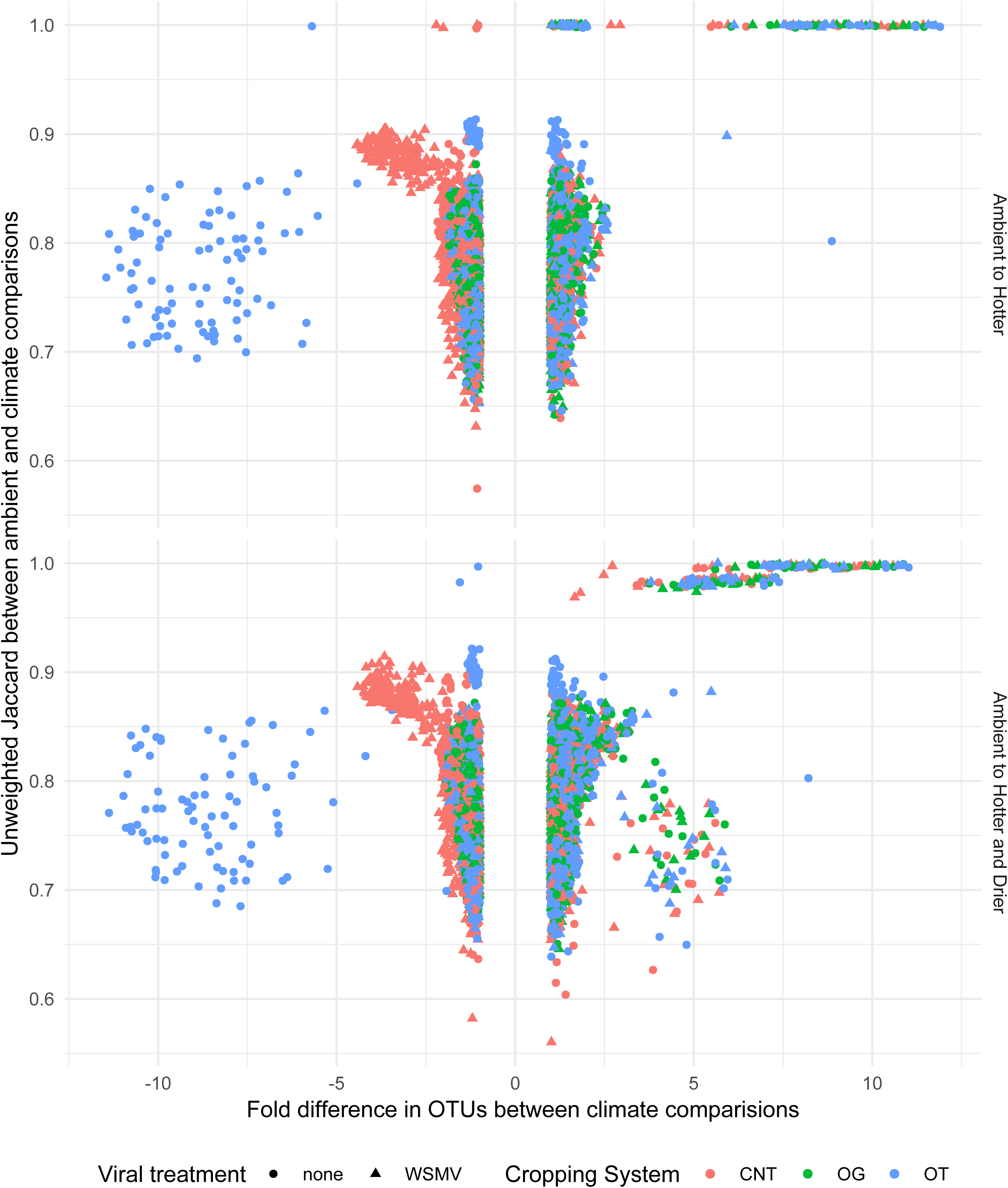
Soil bacterial community similarity against the fold change in number of OTUs in comparing ambient to hotter, and ambient to hotter and drier subplots across the 2016 growing season. Difference in OTUs is measured as fold change, or ratio of the OTU abundance in ambient subplots over the OTU abundance in climate scenario subplots. Viral treatment includes Wheat streak mosaic virus (WSMV) and no-template control (none). Cropping systems include conventional no-till (CNT), organic grazed (OG), and organic tilled (OT).

When comparing bacterial communities in response to climate conditions, cropping system varied how stable bacterial communities remained at different periods in the growing season (Fig 6; Table 3). In May, the bacterial communities in hotter OG samples were less stable compared to the ambient OG samples, as compared to hotter CNT and OT, which were more stable compared to their respective bacterial communities under ambient conditions (Fig 6; Table 3). In July, the bacterial communities in the hotter OT subplots were most stable \, than the hotter CNT or OT subplots and their respective ambient conditions (Fig 6; Table 3). For bacterial communities in hotter and drier subplots as compared to ambient subplots, communities in OT subplots showed the most stability, followed by OG samples, and CNT were least stable (most dissimilar) compared to the respective ambient conditions (Fig 6; Table 3).

**Table 3.**
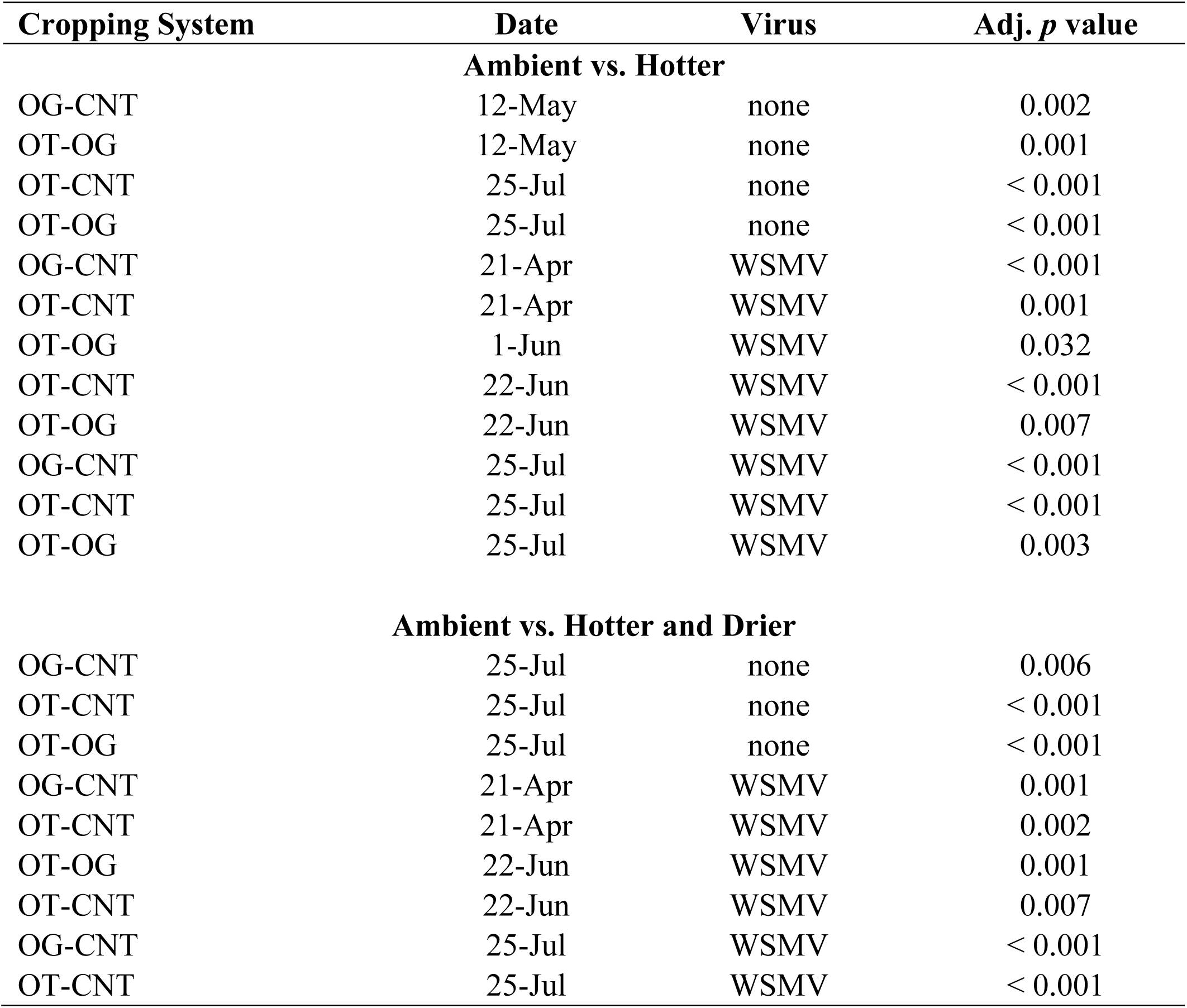
Effect of climate conditions, cropping system, and sampling date on soil bacterial community composition. Unweighted Jaccard was used to calculate bacterial community composition and comparisons were made between ambient and hotter, or ambient and hotter and drier conditions within each sampling date. Comparisons were tested with analysis of variance and P-values adjusted with Tukey’s Honest Significant Differences.

| Cropping System | Date | Virus | Adj. *p* value |
| --- | --- | --- | --- |
| **Ambient vs. Hotter** |  |  |  |
| OG-CNT | 12-May | none | 0.002 |
| OT-OG | 12-May | none | 0.001 |
| OT-CNT | 25-Jul | none | < 0.001 |
| OT-OG | 25-Jul | none | < 0.001 |
| OG-CNT | 21-Apr | WSMV | < 0.001 |
| OT-CNT | 21-Apr | WSMV | 0.001 |
| OT-OG | 1-Jun | WSMV | 0.032 |
| OT-CNT | 22-Jun | WSMV | < 0.001 |
| OT-OG | 22-Jun | WSMV | 0.007 |
| OG-CNT | 25-Jul | WSMV | < 0.001 |
| OT-CNT | 25-Jul | WSMV | < 0.001 |
| OT-OG | 25-Jul | WSMV | 0.003 |
| **Ambient vs. Hotter and Drier** |  |  |  |
| OG-CNT | 25-Jul | none | 0.006 |
| OT-CNT | 25-Jul | none | < 0.001 |
| OT-OG | 25-Jul | none | < 0.001 |
| OG-CNT | 21-Apr | WSMV | 0.001 |
| OT-CNT | 21-Apr | WSMV | 0.002 |
| OT-OG | 22-Jun | WSMV | 0.001 |
| OT-CNT | 22-Jun | WSMV | 0.007 |
| OG-CNT | 25-Jul | WSMV | < 0.001 |
| OT-CNT | 25-Jul | WSMV | < 0.001 |

**Fig 6.**
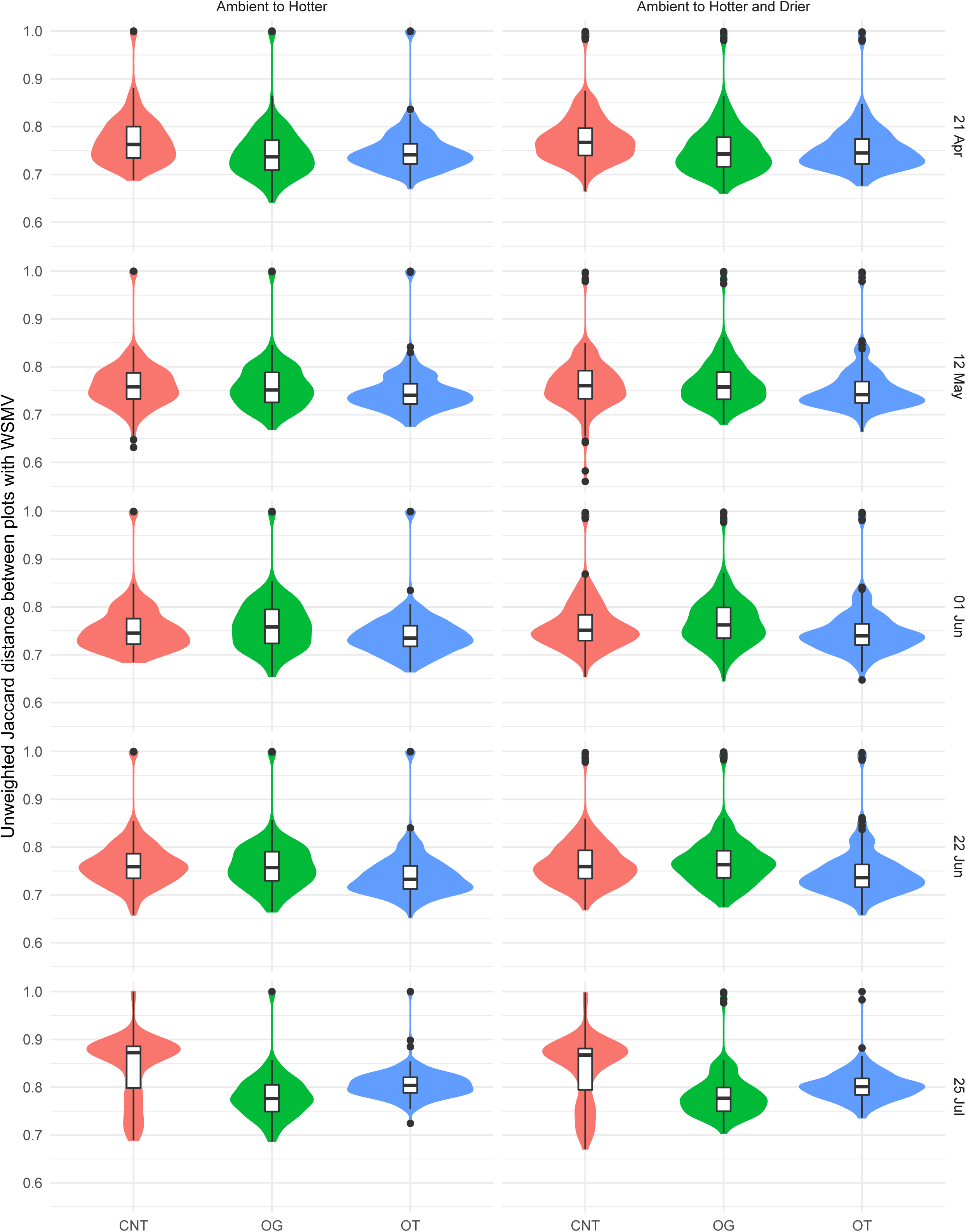
Soil bacterial community similarity between ambient and hotter, and ambient and hotter and drier conditions, subplots from three cropping systems across the 2016 growing season. Plots were not treated with Wheat streak mosaic virus. Significance is provided in Table 4. Cropping systems include conventional no-till (CNT), organic grazed (OG), and organic tilled (OT).

Neither WSMV inoculation nor rate of infection (Table S3) within subplots created a definable bacterial community (random forest, data not shown); although WSMV inoculation was negatively associated with a species of *Cellulomonas*, as well as with Actinobacteria clade 480-2 (Fig S4). However, WSMV inoculation significantly affected soil bacterial community similarity (Table 2). Moreover, there was an interaction of WSMV and climate change (Table 2), which was modulated by cropping system (Fig 7; Table 3).

**Table 4.** PERMANOVA of weed species’ identity and percent coverage (cov) on soil bacterial communities at different times over a growing season. Comparisons were made accounting for repeated measures of subplots, and with replicate blocks as a stratification. Only significant comparisons are shown. Significance was determined as: < 0.001 = ***, 0.001 – 0.009 = **, 0.01 – 0.05 = *, 0.05 - 0.1 = t (trending).

| Weed | F.Model | R2 | p value | Sig |
| --- | --- | --- | --- | --- |
| *Asperugo procumbens* cov Oct 2015 | 1.212 | 0.00474 | 0.047 | * |
| *Bromus tectorum* cov Oct 2015 | 0.819 | 0.0032 | 0.001 | *** |
| *Bromus tectorum* cov Jun 2016 | 1.016 | 0.00397 | 0.005 | ** |
| *Bromus tectorum* biomass late Jun 2016 | 1.1495 | 0.00449 | 0.001 | *** |
| *Capsella bursa-pastoris* cov Oct 2015 | 0.7862 | 0.00307 | 0.035 | * |
| *Capsella bursa-pastoris* cov Apr 2016 | 1.0965 | 0.00429 | 0.001 | *** |
| *Capsella brusa-pastoris* biomass late Jun 2016 | 1.0236 | 0.004 | 0.001 | *** |
| *Chenopodium album* cov Apr 2016 | 1.1424 | 0.00447 | 0.05 | * |
| *Chenopodium album* cov Jun 2016 | 1.0397 | 0.00407 | 0.001 | *** |
| *Cirsium arvense* biomass late Jun 2016 | 0.8499 | 0.00332 | 0.03 | * |
| *Descurainia sofia* cov Oct 2015 | 0.7804 | 0.00305 | 0.003 | ** |
| *Galium aparine* cov Apr 2016 | 0.7757 | 0.00303 | 0.049 | * |
| *Lactuca serriola* biomass late Jun 2016 | 0.9691 | 0.00379 | 0.042 | * |
| *Lamium amplexicaule* cov Oct 2015 | 1.0349 | 0.00405 | 0.047 | * |
| *Malva neglecta* cov Oct 2015 | 1.129 | 0.00441 | 0.019 | * |
| *Malva neglecta* cov Apr 2016 | 0.6295 | 0.00246 | 0.009 | ** |
| *Monolepsis nuttaliana* cov Jun 2016 | 0.9573 | 0.00374 | 0.009 | ** |
| *Poa annua* cov Oct 2015 | 0.8321 | 0.00325 | 0.001 | *** |
| *Poa annua* cov Apr 2016 | 0.7457 | 0.00292 | 0.022 | * |
| *Solanum triflorum* cov Oct 2015 | 0.8559 | 0.00335 | 0.044 | * |
| *Taraxacum officinale* cov Oct 2015 | 0.8523 | 0.00333 | 0.002 | ** |
| *Taraxacum officinale* cov Jun 2016 | 0.7226 | 0.00283 | 0.021 | * |
| *Thlaspi arvense* cov Apr 2016 | 1.3834 | 0.00541 | 0.002 | ** |
| *Thlaspi arvense* cov Jun 2016 | 0.8235 | 0.00322 | 0.047 | * |
| *Tragopogon dubious* biomass late Jun 2016 | 0.799 | 0.00312 | 0.002 | ** |
| *Trifolium pratense* biomass late Jun 2016 | 0.8271 | 0.00323 | 0.001 | *** |
| *Trifolium pratense* cov Apr 2016 | 0.813 | 0.00318 | 0.024 | * |

**Fig 7.**
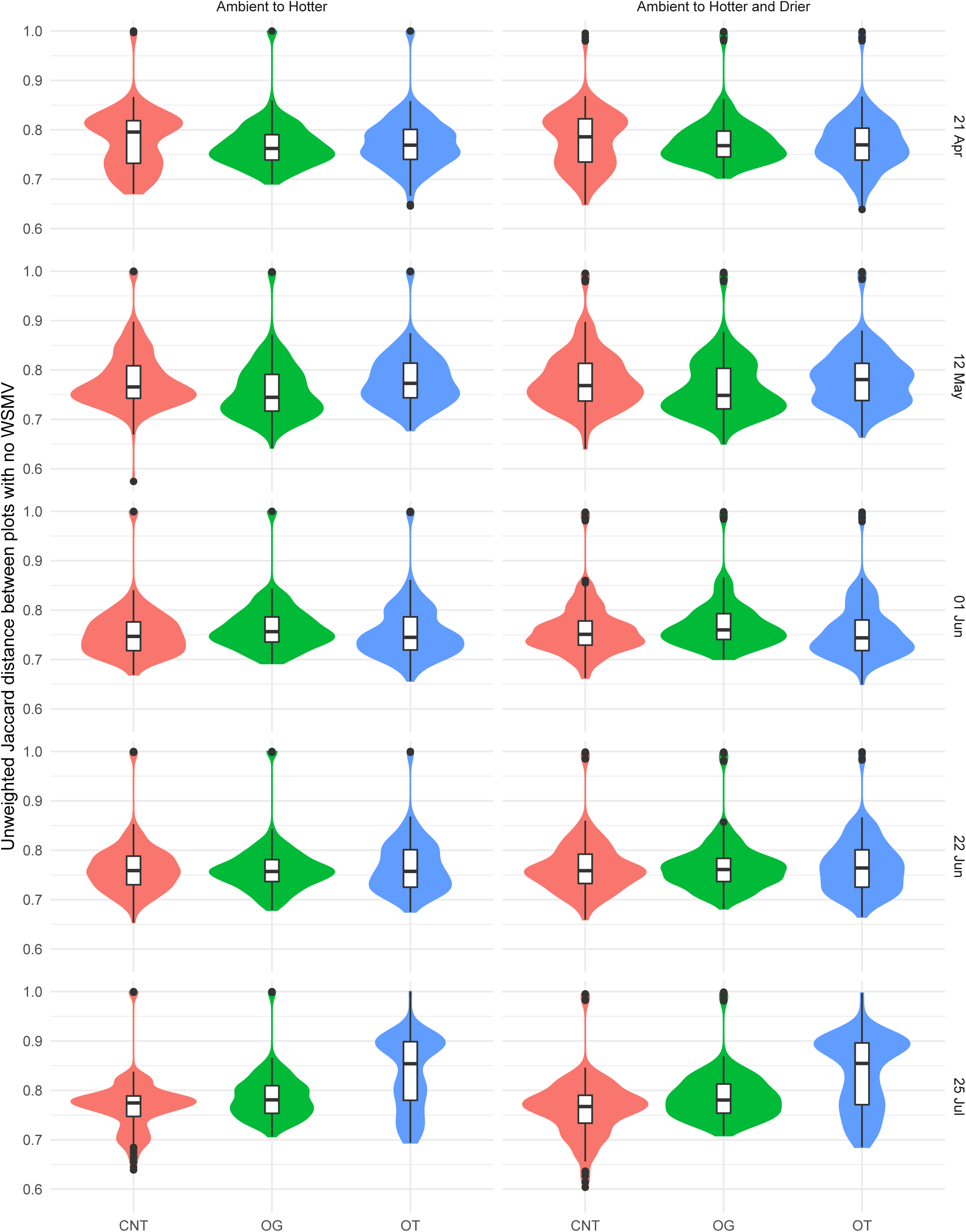
Soil bacterial community similarity between ambient and hotter, and ambient and hotter and drier conditions, in subplots treated with Wheat streak mosaic virus from three cropping systems across the 2016 growing season. Significance is provided in Table 4. Cropping systems include conventional no-till (CNT), organic grazed (OG), and organic tilled (OT).

When comparing the change in bacterial communities between ambient conditions and hotter, or hotter and drier climate conditions, cropping system modulated how stable bacterial communities remained in subplots which had been treated with WSMV. In assessing similarity between bacterial communities between ambient and climate conditions, CNT and OT subplots treated with WSMV were significantly different (ANOVA, *p* < 0.001 Tukey) from their non-infected counterparts, indicating that disease status altered the ability of the community to remain stable (i.e. resistance) under changing climate. However, OG subplots did not differ between WSMV-treated and untreated subplots (ANOVA, *p* > 0.05 Tukey) in terms of the similarity between ambient and climate-conditioned soil.

WSMV made it more difficult for bacterial communities to remain stable under climate conditions, across the growing season, and between cropping systems (Fig 7; Table 3). When comparing ambient to hotter conditions in subplots treated with WSMV, in April, CNT subplots were more stable than OG or OT; in early June, OG was more stable than OT; in late June, CNT and OG were more stable than OT; and in late July, CNT subplots were most stable, followed by OT, and then OG subplots (Fig 7, Table 3). Comparing hotter and drier to ambient conditions in WSMV-treated subplots, in April, CNT subplots were more stable than OG or OT; in late June, CNT and OG were more stable than OT; and in late July, CNT was again more stable than OG or OT (Fig 6; Table 3).

Weed communities in the organic systems were more diverse than the CNT subplots, though OG and CNT had similar relative species abundance (28). Climate conditions had minor impacts on weed communities (28). Weed species’ diversity, as well as percent or biomass from subplots negatively impacted the similarity between ambient and climate-treated subplots across the growing season (Fig S5), including weed diversity (lm, F = 79.153, *p* < 0.001) and percent coverage (F = 26.516, *p* < 0.001) the prior fall on Oct 25, 2015, diversity (F = 25.637, p < 0.001) and coverage (F = 119.78, p < 0.001) early in the growing season on Apr 8, 2016, diversity on 6/14/2016 (F = 68.888, p < 0.001), and weed biomass on June 29, 2016 (F = 30.807, p *<* 0.001).

Individual weed species were associated with membership of the soil bacterial community (Table 4) including *Asperugo procumbens, Bromus tectorum, Capsella brusa-pastoris, Chenopodium album, Cirsium arvense, Descurainia sofia, Galium aparine, Lactuca serriola, Lamium ampleuxicaule, Malva neglecta, Monolepsis nuttaliana, Poa annua, Solanum triflorum, Taraxacum officinale, Thlaspi arvense, Tragopogon dubious*, and *Trifolium pretense.* Of these, three weed species had definable effects on bacterial community structure (Fig S6; *p* < 0.05). Winter annuals, *Bromus tectorum* cover in mid-June (Fig S6A), as well as cover of *Capsella bursa-pastoris* (Fig S6B) and *Descurainia sofia* (Fig S6C) in the previous fall had a predictable impact on the rhizosphere community. *Bromus tectorum* had a U-shaped relationship with bacterial relative abundance, while *C. bursa-pastoris* and *D. sofia* showed more of a positive correlation. *Capsella bursa-pastoris* cover in subplots associated with an increase in *Rubrobacter, Nocardioides, Illumatobacter*, the family-level clade FFCH13075 in the order Solirubrobacteriales, and others (Fig S6B). *Descurainia sofia* coverage of subplots associated with an increase in the KD4-96 clade in the Chloroflexi phylum, FFCH13075, *Blastococcus, Nocadioides, Oryzihumus*, and others (Fig S6C).

## DISCUSSION

This study evaluated the effects of climate conditions, WSMV inoculation, cropping system and associated *in situ* weed communities on wheat soil bacterial communities over the course of a growing season. We hypothesized that 1) bacterial community richness and evenness would be reduced by climate or WSMV infection, 2) cropping systems which promote bacterial richness would be more resistant to alterations from climate and WSMV infection, and 3) more diverse bacterial communities would have a more stable bacterial community membership over the growing season and in response to increased soil temperature, decreased precipitation, and WSMV. In summary, sampling date within the growing season, soil temperature, and soil moisture exerted the greatest effect on soil bacterial communities, followed by cropping system, WSMV infection status, and weed community characteristics.

Changes in precipitation and soil moisture, atmospheric gas concentration, soil salinity, and soil temperature can affect bacterial diversity (19, 20, 48). In particular, soil temperature can be a stronger driver of bacterial diversity and functionality than soil moisture (49). Even after weeks of warmer temperatures, soils microbiota do not appear to develop functional resistance to the heat and maintain stable communities (49), yet soils which experience frequent wet-dry cycles, such as grassland soils, host microbial communities which remain more stable under drought conditions (50). Physical or chemical disturbance can further prevent a stable soil community which is adapted to warmer temperatures from forming (51). In this study, and in accordance previous studies (52, 53), soil temperature was found to be a stronger driver of bacterial species’ diversity and abundance than soil moisture. This may reflect the more complex interaction between plants, microorganisms, and soil conditions, as soil moisture can somewhat stabilize soil temperature (54). Plant foliation, which increases with air temperature, in turn shades soil and can buffer further increases in soil temperature (54), thus better supporting microbial communities.

Cropping systems are known to be associated with particular soil microbial communities (23, 46, 55). In particular, the use of plant- or manure-based fertilizer can increase microbial diversity while chemical-based fertilizers select for acid-tolerant species (46, 56, 57), leading to the trend of organic systems to harbor more diverse microbial communities than conventional (industrial) ones (56, 58, 59). Further, tillage and herbicides reduce microbial diversity (60, 61). Soil microbial α-diversity has been used as a topical application to rescue plants from drought, salt stress, or disease (62–64) and may be used to remediate soils after chemical or physical disruption (65–68). Thus, management practices which promote microbial diversity have the potential to be used as an *in situ* method to moderate the effect of stressors such as climate change, pathogens, or weeds (47, 55).

We previously assessed the impact of these cropping systems over the course of the growing season (23) and observed that under ambient conditions, cropping system did not alter bacterial richness or evenness but did affect β-diversity. In particular, organic tilled subplots contained more putative nitrogen-fixing bacterial genera (23). In the present study, the bacterial community in all cropping systems changed over the course of the growing season, as well as in response to increased soil temperature or decreased soil moisture. However, the interaction between cropping systems and climate conditions, which was not identical across systems. The peak in bacterial richness in CNT and OG systems was delayed in the hotter and hotter and drier conditions as compared to their respective ambient subplots. From observations, wheat in these systems developed more slowly than in OT subplots. The peak in bacterial richness is likely tied to peak growth and development of wheat, when plant-bacterial nutrient exchange is greatest. In OT subplots, the earlier peak and subsequent drop in bacterial richness may be associated with an more advanced growth stage and earlier senescence (13). Cropping system affected the stability of bacterial communities when comparing ambient to climate-treated conditions, with the conventional no-till system remaining more stable than the organic ones. This may reflect the more intense selective pressure exerted by chemical inputs on the community, and the recruitment of a more resilient microbiota.

Cropping system can indirectly alter soil microbial α-diversity via crop disease susceptibility. For example, direct nitrogen fertilization can increase WSMV disease transmission (69). Using livestock grazing to terminate cover crops and control weed residues can reduce wheat mite populations (70), although this has not been shown to reduce virus transmission (71). In the present study, there were interactions between WSMV application and soil moisture, soil temperature, and cropping system X soil moisture, pointing to the importance of multiple concurrent stressors in shaping soil communities. The effect of different cropping systems on viral infection in crops is complex (72) and is largely modulated by the extent of crop diversification, crop residue removal strategy, and pest control (73).

As early successional species, agricultural weeds establish quickly in newly-disturbed soil and sometimes earlier in the growing season than spring or summer crops (74). In climate change scenarios which predict warmer, wetter springs, and higher atmospheric CO_2_, the alteration of the local environmental conditions can give weeds a greater advantage over crops (75). Changing environmental conditions and crop-weed competition may, in turn, alter the soil microbial community, further making conditions less favorable for crop germination, growth, and competitive ability (76). As with all plant species, agricultural weed species associate with particular microbial communities in their rhizosphere (23, 55, 74, 77). It is generally thought that weed diversity in agricultural settings could increase microbial diversity in soil, and potentially increase the functionality and stability of soil microbial communities. In this study site, ambient subplots were previously showed to have weak positive correlations between weed diversity and soil bacterial richness (23). In the present study, weed diversity or biomass did not alter soil bacterial richness or evenness, although bacterial β-diversity was affected, and weed diversity was inversely related to the stability of bacterial communities in response to climate treatment. This may reflect the temporary increases in bacterial richness during periods of weed growth which are not sustained during the hottest part of the season when bacterial communities are more susceptible to temperature and moisture stress.

Environmental conditions or disease status on bacterial communities had interactive effects with cropping system. This has implications for soil bacterial communities and plant performance (78), both within the growing season and in successive plantings, as the legacy of these altered bacterial communities persists (8). As local climates continue to shift, so too will the dynamics of above- and below-ground diversity which will impact food production and the need for more sustainable practices (5, 16, 18).

## MATERIALS AND METHODS

### Experimental Design

This study was conducted in 2015 and 2016 at an agricultural field experiment that was implemented since July 2012 at the Montana State University Fort Ellis Research and Teaching Center, Bozeman, MT (45.652664056 N −110.97249611 W, elevation 1500 m a.s.l.) to test production of three dryland cropping systems using a 5-year crop rotation. The Fort Ellis site is a Blackmore silt loam soil type (a fine-silty, mixed, superactive, frigid Typic Arguistoll) with a consistent ratio of 1 part sand, 2 parts silt, 1 part clay, by weight, at 0 to 4% slopes (79). The monthly air temperature in Bozeman in 2016 was higher than historic maximum and minimums from 1981 – 2010, and the mean monthly precipitation [Table S2, reproduced from (23)] was lower by 18 mm in May, 16 mm in June, and 14 mm in July (80).

The cropping systems at the studied site consisted of 1) conventional no-till system (CNT), in which synthetic inputs were used in the form of fertilizers, herbicides, and fungicides, 2) USDA-certified tilled organic (OT), and 3) USDA-certified organic with grazing (OG), which integrates sheep grazing to terminate cover crops and manage weeds, with the overall goal of reducing tillage intensity in organic production. Chemical inputs utilized in the CNT system included 2,4-D, bromoxynil, dicamba, fluroxypyr, glyphosate, MCPA, pinoxaden, and urea for winter wheat rotations [see Tables 2.7 and 2.8 in (81)]. The organic plots began the organic transition process in July 2012, and completed in 2015. In the OT system, tillage was performed with a chisel plow, tandem disk, or field cultivator, as needed for to control weeds, prepare the seedbed, and to incorporate cover crops and crop residues. Weed control was enhanced with a rotary harrow. In the OG system, targeted sheep grazing was used to reduce tillage intensity for pre-seeding and post-harvest weed control and to terminate the cover crops, with duration and intensity of grazing based on weed biomass (5). Grazing was minimally supplemented with tillage, based on soil conditions and weed pressure. For all systems, seeding was done with a low-disturbance no-till double-disk seeder. Outside of normal farm management activities, soil disturbance and compaction were minimized during sampling procedures. Further details of the management practices, both historical and at the time of experimentation, can be found elsewhere (5, 43, 81).

Each system was replicated three times (i.e. blocks) with cropping systems (75 x 90 m) as the main plots, each of which was further divided into 5 split plots (13 x 90 m), with a 2m fallow buffer in between. Split plots were each following a 5 yr rotation which consisted of: year 1 – safflower (*Carthamus tinctorius* L.) under-sown to yellow sweet clover (*Melilotus oficinalis* (L.) Lam.), year 2 – sweet clover cover crop, year 3 – winter wheat (*Triticum aestivum* L.), year 4 – lentil (*Lens culinaris* Medik.), and year 5 – winter wheat (5).

Within each the year 3 – winter wheat fields, subplots (1 m diameter) were randomly established to assess the impact of climate conditions and disease status on wheat soil bacteria across cropping systems. Two subplots were marked with flags and used as control or ambient climate conditions (ambient), two subplots were enclosed with an open-top chamber (OTC, hotter) made from 18 in high plastic that reflected heat back on the subplot to increase air temperature and soil temperature by 1 - 2° C (82), and two subplots were enclosed with OTCs and partially covered with rain-out shelters (OTC-ROS, hotter and drier) which reduced rainfall by 50% using transparent polyurethane [Fig S7, similar to (83)]. For each of the three climate treatments, one of the subplots was randomly inoculated with WSMV (see below).

### Wheat streak mosaic virus inoculation and data collection

Following previous work (84), prior to the WSMV inoculations, spring wheat (variety Chouteau) was grown in the greenhouse in flat trays (30 x 10 cm), where plants were maintained under a 16-h photoperiod of sunlight supplemented with mercury vapour lamps (165 uE m^-2^ s^-1^) at 10°C/25°C day/night. When the wheat was at Feekes stage 4 - 6, an inoculum of WSMV was created from the ‘Conrad’ isolate line (85). Infected wheat was harvested from the greenhouse and frozen for 1 - 2 days until use. To create the WSMV inoculum, 300 g of infected wheat clippings were ground to reduce particle size using a food processor, then blended with buffer (3.2 L of de-ionized water + 600 ml of 5X PBS, pH 7.2) until smooth. Slurry was filtered through cheesecloth to remove particulate matter which would clog the spray hose, and refrigerated for up to 1 h until use. Immediately prior to use, 2 g carborundum (ground glass) was added per 3.78 L of slurry as an abrasive to slightly injure wheat enough for the virus to infect. Slurry was sprayed onto subplots using an air compressor (275 kilo Pascals) travelling at a rate of 0.5 m/s and sprayed at a height of 20 cm above the canopy. Control subplots were sprayed with water in which 2 g carborundum was added per 3.78 L (no-template control). Spraying occurred the last week of April, one week after the first soil sampling date (April 21) and two weeks prior to the second sampling date (May 12).

Infection of WSMV in subplots was evaluated in July by using an indirect ELISA, with 10 leaves sampled from each subplot and assessed separately (Ito et al. 2012). Within a plate, every 10th well contained a negative control (i.e., sample from healthy wheat plant) to reduce potential bias in values of optical density caused by position of samples. The mean and standard deviation of the negative control on each plate were calculated. Samples above three standard deviations were considered infected with WSMV (Miller et al. 2014). ELISA results are provided in Table S1.

### Crop and weed evaluations

Percent coverage of weeds in subplots was assessed visually in October 2015, April 2016, and June 2016. Aboveground biomass of all weed species within sampled areas was harvested by hand in late June 2016. Within each 0.75 m^2^ subplot, weed biomass was cut at ground level and separated by species. The individual biomass of each species was dried for 2 weeks at 55° C, and weighed (28). Wheat biomass was harvested from sampled areas by hand on July 25, 2016, once the crop had completely senesced and ripened. The two center rows (75 cm each) of wheat in the subplot were harvested, for a total of 1.5 row meters. All the aboveground biomass was harvested, dried for 1 week at 55° C and threshed to determine biomass and grain yields (28)

### Soil assessment

Soil moisture was measured weekly using gypsum blocks buried at 5 cm below ground (86). Soil temperature was measured with buried iButtons (Maxim Integrated), with data obtained every four hours between April 14, 2016 (one week prior to the first sampling) and July 25, 2016 (final sampling date). In each subplot, three cores were taken from around wheat plants to a depth of 15 cm, then homogenized into one composite sample, which was used for bacterial community sampling (stored at −20°C) and nutrient analysis (stored at 4°C). Soil cores were obtained from all 54 subplots at five time-points over the growing season: April 21 before the WSMV inoculations were applied; May 12, one week post-WSMV infection; June 1, three weeks post-WSMV infection; June 22, six weeks post-WSMV infection; and July 25^th^, 10 weeks post-WSMV infection and immediately prior to wheat harvesting. Additional soil was collected at wheat harvest for nutrient analysis, presented in Table S3 (Agvise Laboratories, Northwood, North Dakota, US).

DNA extraction from soil samples, library preparation, sequencing, and sequence analysis protocols were as previously described (23). Illumina MiSeq (Montana State University, Bozeman, MT) was used to sequence the V3-V4 region of the 16S rRNA gene, using primers 341F (5’-CCTACGGGAGGCAGCAG-3’) and 806R (5’-GGACTACHVGGGTWTCTAAT-3’) (87). Sequencing output data can be found in the Sequence Read Archive (SRA) at NCBI under BioProject PRJNA383161 (https://www.ncbi.nlm.nih.gov/bioproject/?term=PRJNA383161).

Linear mixed effects models and distance based redundancy models (vegan) (88), random forest with permutation (89, 90), PERMANOVA (adonis) (91), and ggplot2 (92) were used in the R statistical package (93). Linear mixed models used cropping system, soil moisture and soil temperature on the day of sampling, and WSMV application as random effects. Sampling date and subplot identity nested with block were included to control for repeated sampling. Some variables were aliased in the distance-based redundancy analysis and therefore were removed from the model: *Capsella brusa-pastori*s, *Cirsium arvense, Galium aparine*, and *Tragopogon dubious* biomass on June 29, 2016; *Chenopodium album, Lamium ampleuxicaule, Malva neglecta, Poa annua*, and *Solanum triflorum* coverage October 25, 2015; *C. arvense, T. dubious*, and *Trifolium pretense* coverage April 8, 2016; and *Chenopodium album* and *Tragopogon dubious* coverage June 14, 2016. Plant coverage has been shown to have a linear correlation with plant aboveground biomass (94, 95) and, weed senescence may negate the effect on soil microorganisms (95). Thus, as coverage was measured at multiple timepoints but biomass only once, coverage was used as a more accurate measure of the weed-soil microbe relationship with respect to sampling date when coverage and biomass were both significant.

Random forest was performed with 500 trees and 100 permutations. Replicate block did not affect numerical diversity and was included as a random effect in those models, but did affect bacterial communities when comparing ambient systems (23), and was included as a fixed effect in those models. Unweighted Jaccard similarity was used to determine effect of factors on community structure, and tested with PERMANOVA (adonis), with replicate block as a stratification. When comparing climate to ambient conditions, we utilized analysis of variance (ANOVA) and Tukey’s Honest Significant Differences to assess the variables determining soil bacterial communities. The comparison and visualization of ambient to climate conditions was based on R code developed by Drs. Ashkaan Fahimipour and Roo Vandegrift.

## ACKNOWLEDGEMENTS

The authors would like to thank Kyla Crisp, Madison Nixon, Tessa Scott, Rachel Flowers, Ali Thornton, and Lazaro Vinola for their assistance maintaining the plots and collecting samples; Dr. Mary Burrows and Everett Owen for assistance producing Wheat streak virus; Devon Ragen for sheep maintenance; Drs. Pat Hatfield and Perry Miller for farm administration; Sarah Olivo for DNA sequencing; and Genna Shaia for provided some literature review assistance. This work was supported by the USDA NIFA Organic Transitions (ORG) program (Grant MONB00128), the Montana Agricultural Experiment Station (project MONB00113), and the National Institute of General Medical Sciences of the National Institutes of Health (NIH-NIGMS; award number P20GM103474).

